# Decoding the temporal dynamics of affective scene processing

**DOI:** 10.1101/2022.01.27.478067

**Authors:** Ke Bo, Lihan Cui, Siyang Yin, Zhenhong Hu, Xiangfei Hong, Sungkean Kim, Andreas Keil, Mingzhou Ding

## Abstract

Natural images containing affective scenes are used extensively to investigate the neural mechanisms of visual emotion processing. Functional fMRI studies have shown that these images activate a large-scale distributed brain network that encompasses areas in visual, temporal, and frontal cortices. The underlying spatial and temporal dynamics among these network structures, however, remain to be characterized. We recorded simultaneous EEG-fMRI data while participants passively viewed affective images from the International Affective Picture System (IAPS). Applying multivariate pattern analysis to decode EEG data, and representational similarity analysis to fuse EEG data with simultaneously recorded fMRI data, we found that: (1) ~100 ms after picture onset, perceptual processing of complex visual scenes began in early visual cortex, proceeding to ventral visual cortex at ~160 ms, (2) between ~200 and ~300 ms (pleasant pictures: ~200 ms; unpleasant pictures: ~260 ms), affect-specific neural representations began to form, supported mainly by areas in occipital and temporal cortices, and (3) affect-specific neural representations, lasting up to ~2 s, were stable and exhibited temporally generalizable activity patterns. These results suggest that affective scene representations in the brain are formed in a valence-dependent manner and are sustained by recurrent neural interactions among distributed brain areas.

## INTRODUCTION

The visual system detects and evaluates threats and opportunities in complex visual environments to facilitate the organism’s survival. In humans, to investigate the underlying neural mechanisms, we record fMRI and/or EEG data from observers viewing depictions of naturalistic scenes varying in affective content. A large body of previous fMRI work has shown that viewing emotionally engaging pictures, compared to neutral ones, heightens blood flow in limbic, frontoparietal, and higher-order visual structures (Lang et al., 1998; Phan et al., 2002; Liu et al., 2012; Bradley et al., 2015). Applying MVPA and functional connectivity techniques to fMRI data, we further reported that affective content can be decoded from voxel patterns across the entire visual hierarchy, including early retinotopic visual cortex, and that the anterior emotion-modulating structures such as the amygdala and the prefrontal cortex are the likely sources of these affective signals via the mechanism of reentry (Bo et al., 2021).

Temporal dynamics of affective scene processing remains to be better elucidated. The event-related potential (ERP), an index of average neural mass activity with millisecond temporal resolution, has been the main method for characterizing the temporal aspects of affective scene perception (Cuthbert et al., 2000; Keil et al., 2002; Hajcak et al., 2009). Univariate ERPs are sensitive to local neural processes but do not reflect the contributions of multiple neural processes taking place in distributed brain regions underlying affective scene perception. The advent of the multivariate decoding approach has begun to expand the potential of the ERPs (Bae and Luck 2019; Sutterer et al., 2021). By going beyond univariate evaluations of condition differences, these multivariate pattern analyses (MVPA) take into account voltage topographies reflecting distributed neural activities, and uncover the discriminability of experimental conditions not possible with the univariate ERP method. The MVPA method can even be applied to single-trial EEG data. By going beyond mean voltages, the decoding algorithms can examine differences in single-trial EEG activity patterns across all sensors, which further complements the ERP method (Grootswagers et al., 2017; Contini et al., 2017). Conceptually, the presence of decodable information in neural patterns has been taken to index differences in neural representations (Norman et al., 2006). Thus, in the context of EEG/ERP data, the time course of decoder performance may inform on how neural representations linked to a given condition or stimulus form and evolve over time (Cauchoix et al 2014; Wolff et al., 2015; Dima et al., 2018).

The first question we considered was how long it takes for the neural representations of affective scenes to form. For non-affective images containing objects such as faces, houses or scenes, past work has shown that the neural responses become decodable as early as ~100 ms after stimulus onset (Cichy et al., 2014; Cauchoix et al., 2014). This latency reflects the initial detection and categorization of stereotypical visual features associated with different objects in early visual cortex (Nakamura et al 1997; Di Russo et al. 2002). For complex scenes varying in affective content, however, there are no stereotypical visual features that unambiguously separate different affective categories (e.g., unpleasant scenes vs neutral scenes). Accordingly, univariate ERP studies have reported robust voltage differences between emotional and neutral content at relatively later onset times, e.g., ~170-280 ms at the level of the early posterior negativity (Schupp et al., 2006; Foti et al., 2009) and ~300 ms at the level of the late positive potential (LPP) (Cuthbert et al., 2000, Lang and Bradley, 2010; Liu et al., 2012; Sabatinelli et al., 2013). We sought to further examine these issues by applying multimodal neuroimaging and the MVPA methodology. It is expected that perceptual processing of affective scenes would begin ~100 ms following picture onset whereas affect-specific neural representations would emerge between ~150 ms and ~300 ms.

A related question is whether there are systematic timing differences in the formation of neural representations of affective scenes differing in emotional content. Specifically, it has been debated to what extent pleasant versus unpleasant contents emerge over different temporal intervals (e.g. Oya et al., 2002). The negativity bias idea suggests that aversive information receives prioritized processing in the brain and predicts that scenes containing unpleasant elements evoke faster and stronger responses compared to scenes containing pleasant or neutral elements. The ERP results to date have been equivocal (Carretié et al., 2001; Huang and Luo, 2006; Franken et al., 2008). An alternative idea is that the timing of emotional representation formation depends on the specific content of the images (e.g., erotic within the pleasant category vs mutilated bodies within the unpleasant category) rather than on the broader semantic categories such as unpleasant scenes and pleasant scenes (Weinberg and Hajcak, 2010). We sought to test these ideas by applying the MVPA approach to decode scalp topographies of single-trial EEG data.

How do neural representations of affective scenes, after their formation, evolve over time? For non-affective images, the neural responses are found to be transient, with the processing locus evolving dynamically from one brain structure to another (Carlson et al., 2013; Cichy et al., 2014; Kaiser et al., 2016). For affective images, in contrast, the enhanced LPP, a major ERP index of affective processing, is persistent, lasting up to several seconds, and supported by distributed brain regions including the visual cortex as well as frontal structures, suggesting sustained representations. To test whether neural representations of affective scenes are dynamic or sustained, we applied a MVPA method called the generalization across time (GAT) (King and Dehaene, 2014), in which the MVPA classifier is trained on data at one time point and tested on data from all time points. The resulting temporal generalization matrix, when plotted on the plane spanned by the training time and the testing time, can be used to visualize the temporal stability of neural representations. For a dynamically evolving neural representation, high decoding accuracy will be concentrated along the diagonal in the plane, namely, the classifier trained at one time point can only be used to decode data from the same time point but not data from other time points. For a stable or sustained neural representation, on the other hand, high decoding accuracy extends away from the diagonal line, indicating that the classifier trained at one time point can be used to decode data from other time points. It is expected that the neural representations of affective scenes are sustained rather than dynamic with the visual cortex playing an important role in the sustained representation.

We recorded simultaneous EEG-fMRI data from participants viewing affective images from the International Affective Picture System (IAPS) (Lang et al., 1997) (see Figure 1A for the paradigm). MVPA was applied to EEG data to assess the formation of affective scene representations in the brain and their stability. EEG and fMRI data were integrated to assess the role of visual cortex in the large-scale recurrent network interactions underlying the sustained representation of affective scenes. Fusing EEG and fMRI data via representation similarity analysis (RSA), we were able to further test the timing of perceptual processing of affective scenes in areas along the visual pathway and compare that with the formation time of affect-specific representations. The analysis pipeline is shown in Figure 1B.

**Figure 1.**
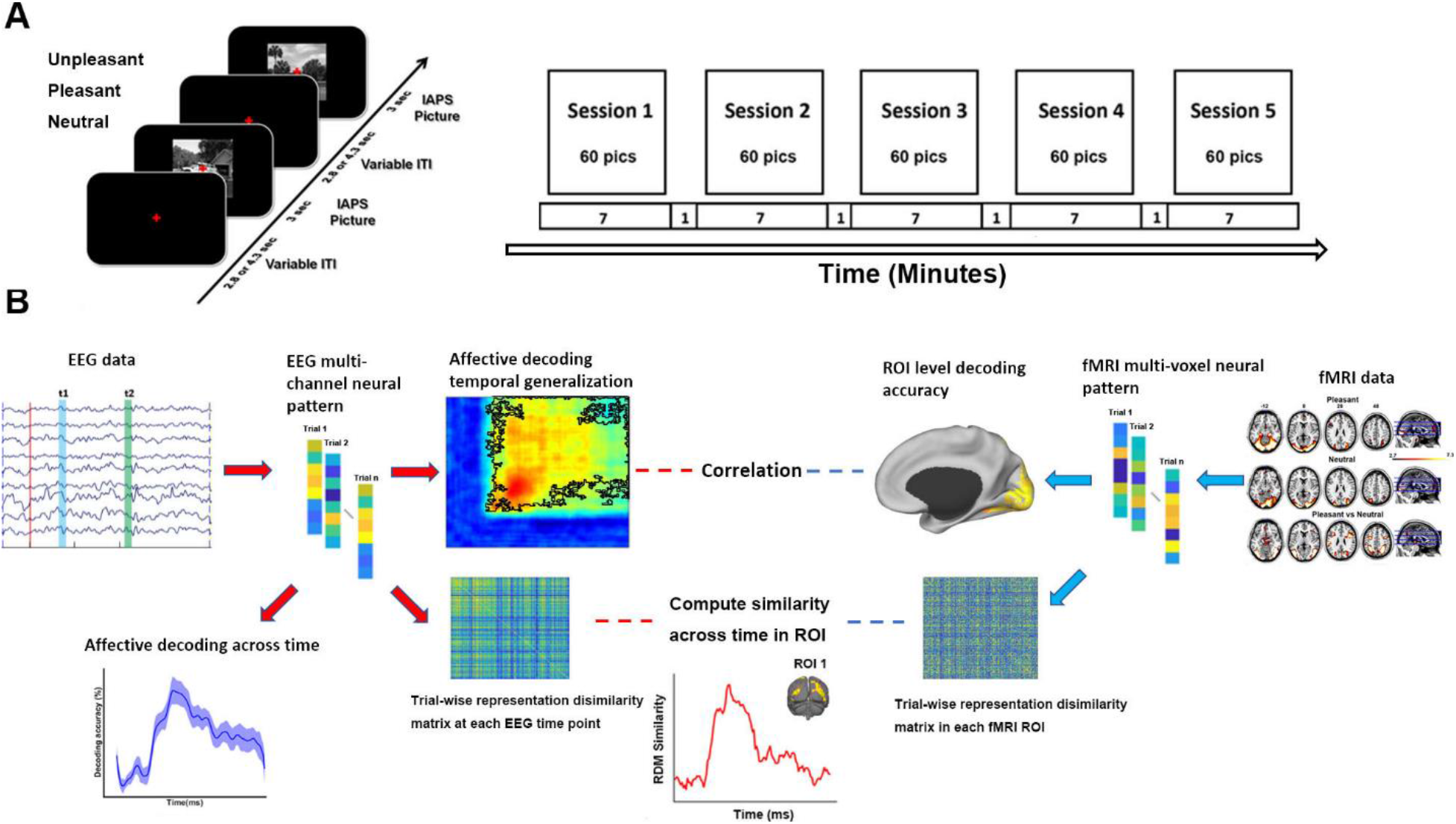
Experiment and data analysis pipeline. **A)** Affective picture viewing paradigm. Each recording session lasts seven minutes. 60 IAPS pictures including 20 pleasant, 20 unpleasant and 20 neutral pictures were presented in each session in random order. Each picture was presented at the center of screen for 3 seconds and followed by a fixation period (2.8 or 4.3 seconds). Participants were required to fixate the red cross at the center of the screen throughout the session while simultaneous EEG-fMRI was recorded. **B)** Analysis pipeline with the methods used at different steps of the analysis (see text for more details).

## RESULTS

### Affect-specific neural representations: Onset time

We decoded multivariate EEG patterns evoked by pleasant, unpleasant, and neutral affective scenes and obtained the decoding accuracy time courses for pleasant-vs-neutral and unpleasant-vs-neutral. As shown in Figure 2A, for pleasant vs neutral, above-chance level decoding began ~200 ms after stimulus onset, whereas for unpleasant vs neutral, the onset time of above-chance decoding was ~260 ms. Using a bootstrap procedure, the distributions of the onset times were obtained and shown in Figure 2B, where the difference between the two distributions was evident, with pleasant-specific representations forming significantly earlier than that of unpleasant-specific representations (ks value = 0.87, effect size = 1.49, two-sample Kolmogorov-Smirnov test). To examine the contribution of different electrodes to the decoding performance, Figure 2C shows the classifier weight maps at the indicated times. These weight maps suggested that neural activities that contributed to classifier performance was mainly located in occipital-temporal channels, in agreement with prior studies using fMRI where enhanced BOLD activity by affective scenes was observed in visual cortex and temporal structures (Sabatinelli et al., 2006; Sabatinelli et al., 2013; Bo et al., 2021).

**Figure 2.**
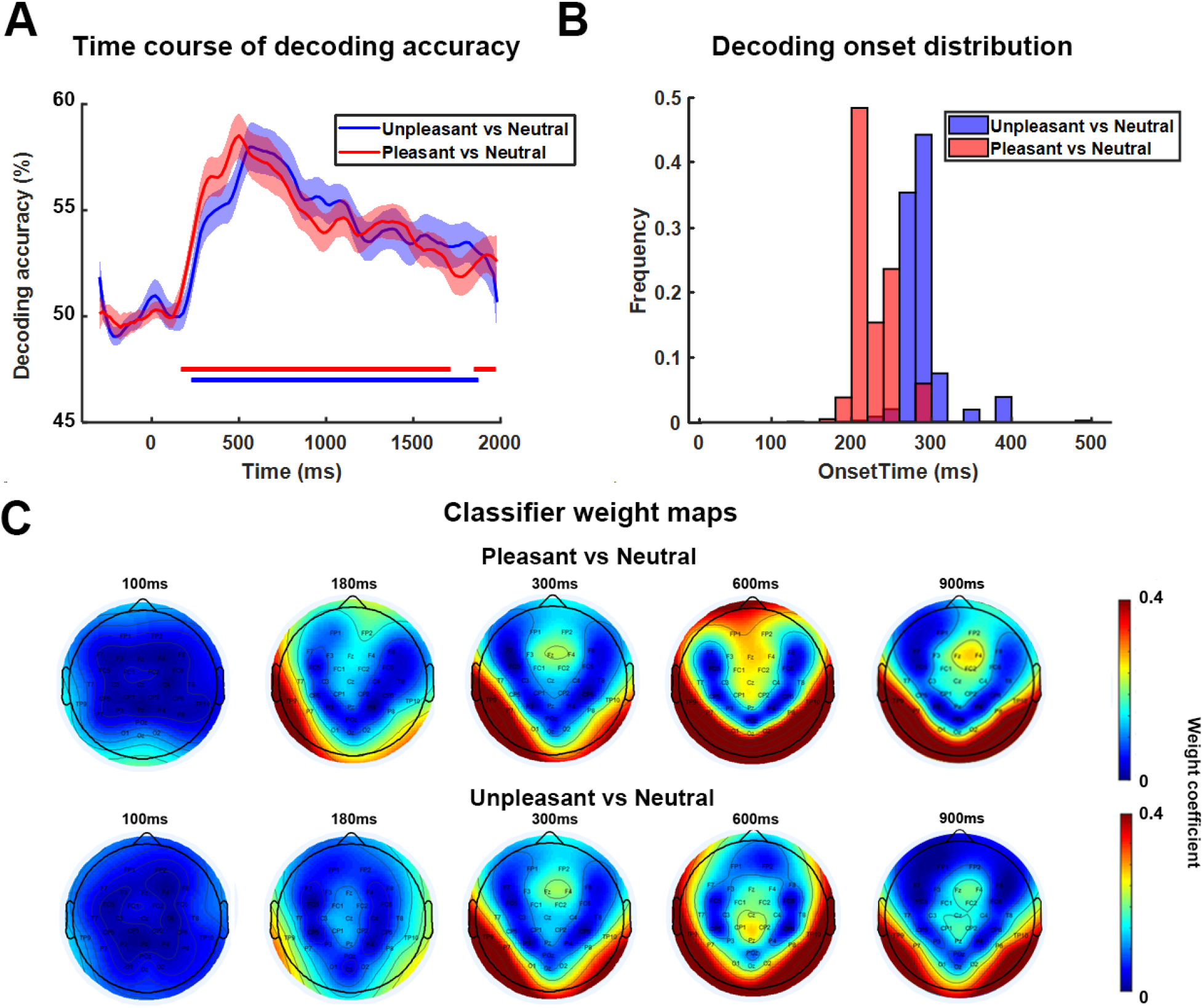
Decoding EEG data between affective and neutral scenes across time. **A)** Decoding accuracy time courses. **B)** Bootstrap distributions of above-chance decoding onset times. Subjects are randomly selected with replacement and onset time was computed for each bootstrap resample (a total of 1000 resamples were considered). **C)** Weight maps showing the contribution of different channels to decoding performance at different times.

Further dividing the scenes into 6 subcategories: erotic couple, happy people, mutilation body/disgust, attack, nature scene/adventure, and neutral people, we decoded multivariate EEG patterns evoked by these subcategories of images. Against neutral people, the onset times of above-chance decoding for erotic couple, attack, and mutilation body/disgust were ~180 ms, ~280 ms, and ~300 ms, respectively, with happy people not significantly decoded from neutral people. The onset times were significantly different between erotic couple and attack with erotic couple being earlier (ks value = 0.81, effect size = 2.1), and between erotic couple and mutilation body/disgust with erotic couple being earlier (ks value=0.92, effect size = 2.3). The onset times between attack and mutilated body/disgust were only weakly different with attack being earlier (ks value = 0.35, effect size = 0.34). Against natural scenes, the onset times of above-chance level decoding for erotic couple, attack, and mutilation body/disgust were ~240 ms, ~300 ms, and ~300 ms, respectively, with happy people not significantly decoded from natural scenes. The onset times were significantly different between erotic couple and attack with erotic couple being earlier (ks value = 0.7, effect size = 1.3) and between erotic and mutilation body/disgust with erotic couple being earlier (ks value = 0.87, effect size = 1.33); the onset timings were not significantly different between attack and mutilation body/disgust (ks value = 0.25, effect size = 0.25). Combining these data, for subcategories of affective scenes, the formation time of affect-specific neural representations appear to follow the temporal sequence: erotic couple → attack → mutilation body/disgust.

### Affect-specific neural representations: Temporal stability

How do affect-specific neural representations, once formed, evolve over time? A serial processing model, in which neural processing progresses from one brain region to the next, would predict that the representations will evolve dynamically, resulting in a temporal generalization matrix as schematically shown in Figure 3A Left. In contrast, a recurrent processing model, in which the representations are undergirded by the recurrent interactions among different brain regions, would predict sustained neural representations, resulting in a temporal generalization matrix as schematically shown in Figure 3A Right. We applied the generalization across time (GAT) method to test these possibilities. A classifier was trained on data recorded at time *t_y_* and tested on data at time *t_x_*. The decoding accuracy is then displayed as a color-coded two-dimensional function (called the temporal generalization matrix) on the plane spanned by *t_x_* and *t_y_*. As can be seen in Figure 3B, a stable neural representation emerged ~200 ms after picture onset and remained stable as late as 2000 ms post stimulus onset, with the peak decoding accuracy occurring within the time interval 300 ms-800 ms. Although the decoding accuracy decreased after the peak time, it remained significantly above chance, as shown by the large area within the black contour. These results demonstrate that the affect-specific neural representations of affective scenes, whether pleasant or unpleasant, are stable and sustained over extended periods of time, suggesting recurrent interactions in the engaged neural circuits. Repeating the same temporal generalization analysis for emotional subcategories, as shown in Figure 4, we observed similar stable neural representations for each emotion subcategory, consistent with Figure 3, suggesting that recurrent interactions also characterize the temporal evolution of the affect-specific representations of emotion subcategories.

**Figure 3.**
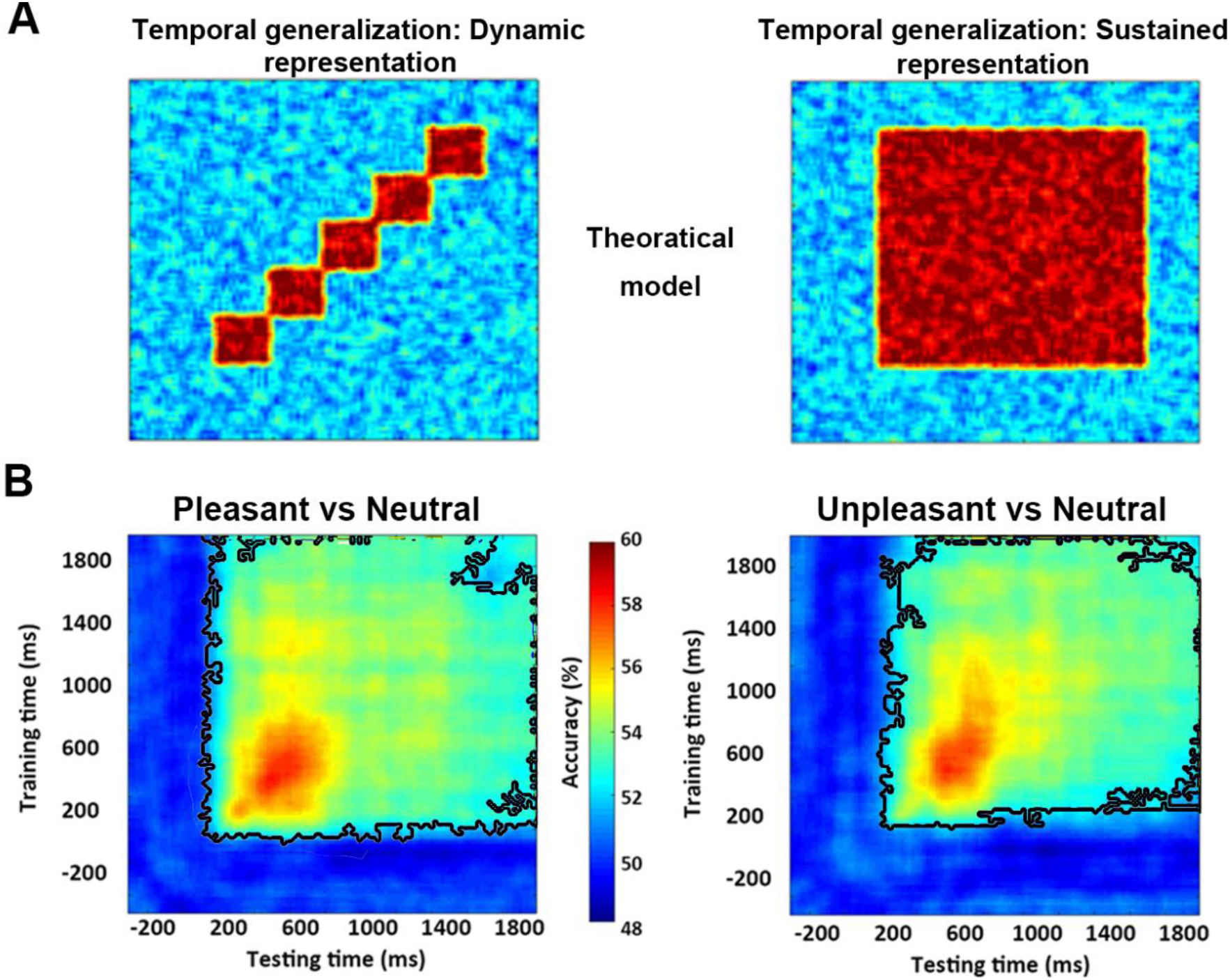
Temporal stability analysis. Classifier trained at each time point was tested on other time points in the time series. The decoding accuracy at a point on this plane reflects the performance at time *t_x_* of the classifier trained at time *t_y_*. **A)** Schematic temporal generalizations of dynamic or transient (Left) vs sustained or stable (Right) neural representations. **B)** Temporal generalization for decoding between pleasant vs neutral (Left) and between unpleasant vs neutral (Right). Black contours outline the statistically significant pixels (p<0.05, FDR)

**Figure 4.**
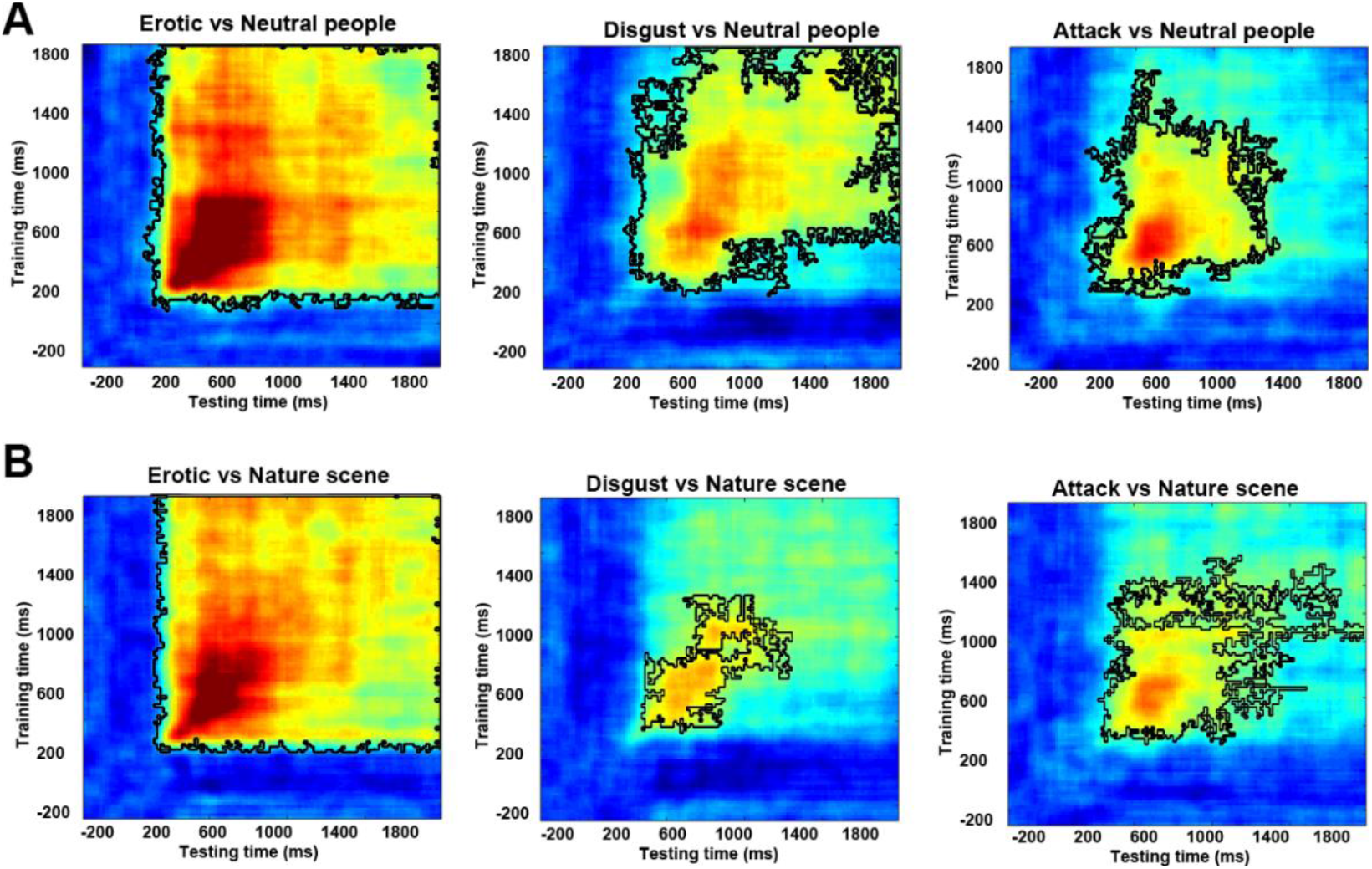
Temporal generalization of affect-specific neural representations of subcategories of affective scenes. **A)** Decoding emotion subcategories against neutral people. **B)** Decoding emotion subcategories against natural scenes. Black contours outline the statistically significant pixels (p<0.05, FDR).

### Visual cortical contributions to sustained affective representations

Weight maps in Figure 2 suggest that occipital and temporal structures are the main neural substrate underlying affect-specific neural representations, which is in line with previous studies showing patterns of visual cortex activity encoding rich, category-specific emotion representations (Kragel et al., 2019; Bo et al., 2021). Whether these structures participate in the recurrent interactions that give rise to sustained neural representations of affective scenes was the question we considered next. Previous work has shown that cognitive operations such as attention, working memory, and decision-making are characterized by sustained neural representations, assessed by temporal generalization, in which sensory cortex is an essential node in the recurrent network (Büchel and Friston, 1997; Gazzaley et al., 2004; Wimmer et al., 2015). We tested whether the same holds true in affective scene processing. Figure 5A shows above-chance fMRI decoding accuracy for pleasant vs neutral (p<0.001) and unpleasant vs neutral (p<0.001) in visual cortex. If the visual cortex is engaged in recurrent interactions with anterior emotion structures, then the stronger and longer the recurrent interactions persist, the stronger and more distinctive the affective representations in the visual cortex. We quantified the strength of the temporal generalization matrix by averaging the decoding accuracy in the black contour (see Figure 3B) and correlated this strength with the fMRI decoding accuracy in visual cortex. As shown in Figure 5B, for unpleasant vs neutral decoding, there was a significant correlation between fMRI decoding accuracy in visual cortex and the strength of temporal generalization (R=0.66, p=0.0008), whereas for pleasant vs neutral decoding, the correlation is not as strong but is still marginally significant (R=0.32, p=0.07). Dividing subjects into high and low decoding accuracy group based on their fMRI decoding accuracies in the visual cortex, the corresponding temporal generalization for each group is shown in Figure 5C, where it is again intuitively clear that temporal generalization is stronger in subjects with higher decoding accuracy in the visual cortex. Statistically, the strength of temporal generalization for unpleasant vs neutral was significantly larger in the high decoding accuracy group (p=0.01) than the low accuracy group; the same was also observed for pleasant vs neutral but the statistical effect is again weaker (p=0.065).

**Figure 5.**
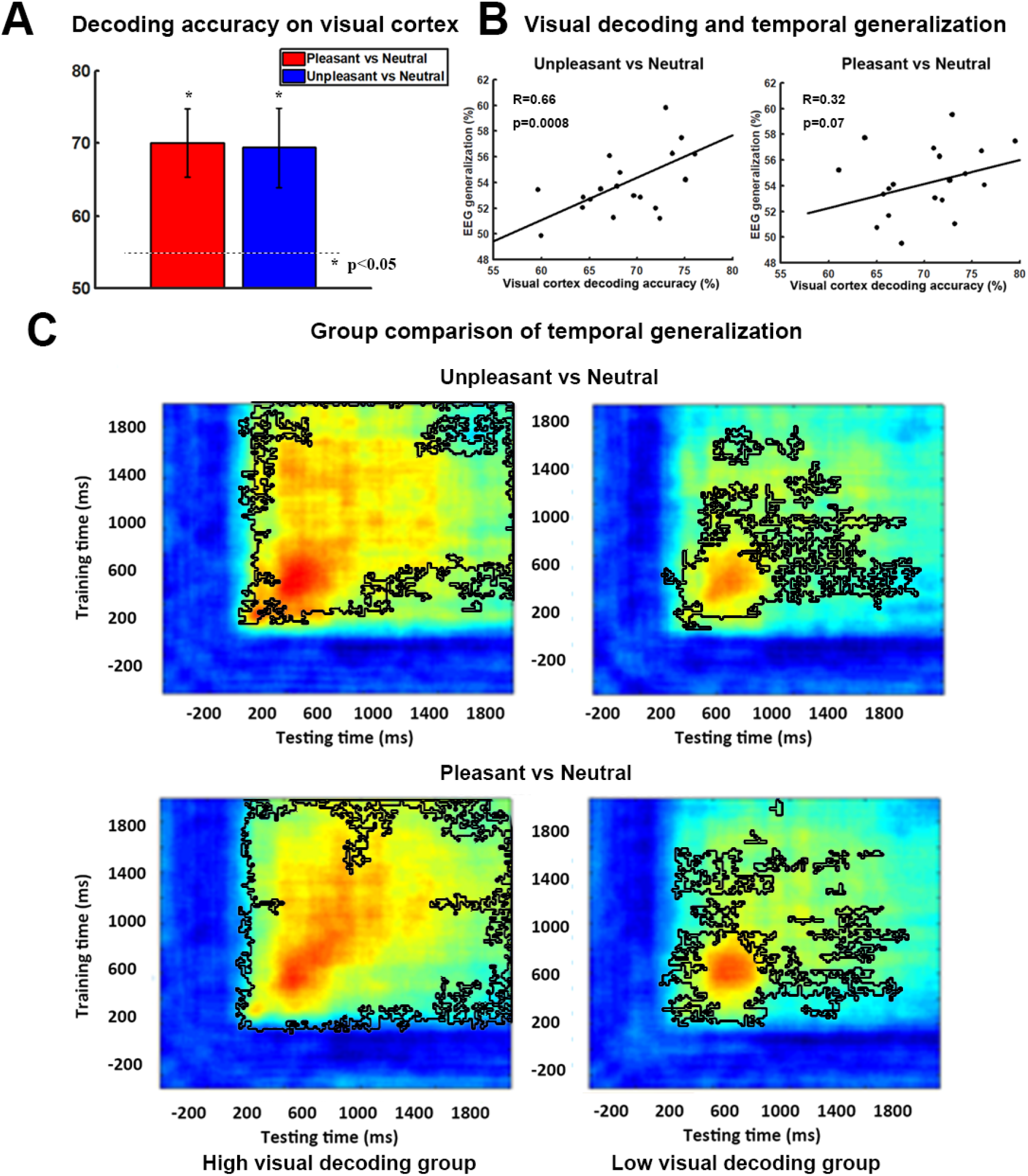
Visual cortical contribution to stable representations of affect. **A)** fMRI decoding accuracy in visual cortex. P<0.05 threshold indicated by the dashed line. **B)** Correlation between strength of EEG temporal generalization and fMRI decoding accuracy in visual cortex. **C)** Subjects are divided into two groups according to their fMRI decoding accuracy in visual cortex. Temporal generalization for pleasant vs neutral and unpleasant vs neutral was shown for each group (high accuracy group on the Left vs low accuracy group on the Right). Black contours outline the statistically significant pixels (p<0.05, FDR).

### Timing of perceptual processing of affective scenes

Past work has found that perceptual processing of simple visual objects begin ~100 ms after image onset in visual cortex (Cichy et al., 2016). This time is earlier than the formation time of affect-specific neural representations (~200 ms). Since the present study used complex visual scenes, it would be helpful to obtain information on the timing of perceptual processing of these images and to provide a reference for comparison. We fused simultaneous EEG-fMRI data using representational similarity analysis (RSA) (Cichy et al., 2016; Cichy and Teng, 2017) and compute the time at which visual processing of IAPS images began in visual cortex. Visual cortex was subdivided into early, ventral, and dorsal parts (see Methods). Their anatomical locations are shown in Figure 6A. We found that shared variance between EEG recorded on the scalp and fMRI recorded from early visual cortex (EVC), ventral visual cortex (VVC), and dorsal visual cortex (DVC) began to exceed statistical significance level at ~100 ms, ~160 ms, and ~360 ms post picture onset, respectively, and remained significant until ~1800 ms; see Figure 6B. These onset times are significantly different according to the KS test applied to bootstrap generated onset time distributions: EVC<VVC (ks value = 0.48 and effect size = 0.49), VVC<DVC (ks value = 0.87 and effect size = 1.29), and EVC<DVC (ks test = 0.95, effect size =1.93); see Figure 6C.

**Figure 6.**
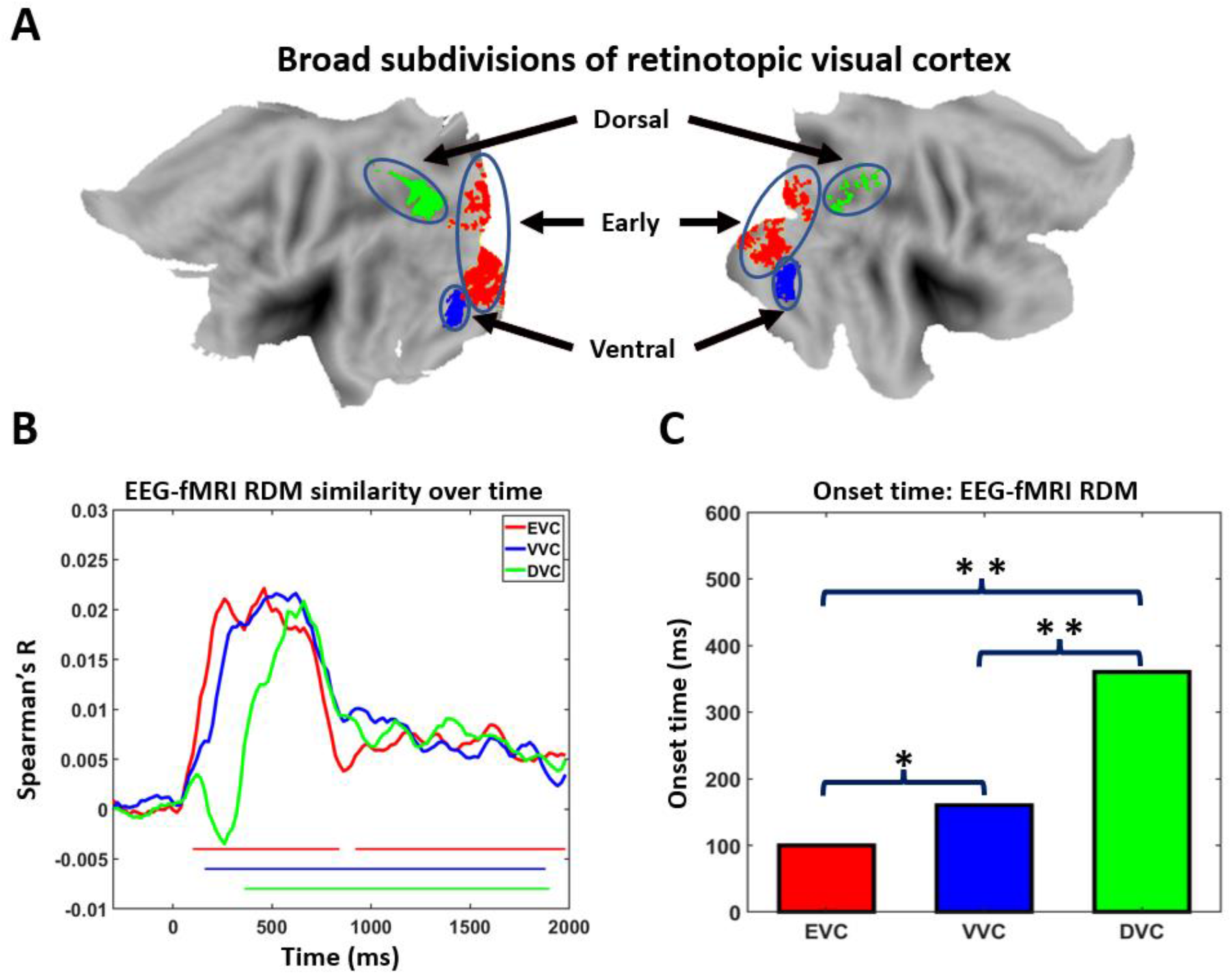
Time course of perceptual processing of complex natural scenes in visual cortex. **A)** Regions of interest (ROIs): early visual cortex (EVC), ventral visual cortex (VVC), and dorsal visual cortex (DVC). **B)** RSA analysis. Left: Similarity between EEG RDM and fMRI RDM across time for the three ROIs. Similarity larger than five baseline standard deviations for at least 5 consecutive time points are marked as statistically significant. **C)** Onset time of similarity curve for each ROI. * Medium effect size. ** Large effect size.

## DISCUSSION

We investigated the temporal dynamics of affective scene processing and reported four main observations. First, EEG patterns evoked by both pleasant and unpleasant scenes were distinct from those evoked by neutral scenes, with above-chance decoding occurring ~200 ms post image onset. The formation of pleasant-specific neural representations led that of unpleasant-specific neural representations by about 60 ms (~200 ms vs ~260 ms); the peak decoding accuracies were about the same (59% vs 58%). Second, dividing affective scenes into six subcategories, the onset of above-chance decoding between affective and neutral scenes followed the sequence: erotic couple (~210 ms)→attack (~ 290 ms)→mutilation body/disgust (~300 ms), suggesting that the speed at which neural representations form depends on specific picture content. Third, for both pleasant and unpleasant scenes, the neural representations were sustained rather than transient, and the stability of the representations was associated with the fMRI decoding accuracy in the visual cortex, suggesting an active role of visual cortex in the recurrent neural network that supports the representations. Fourth, applying RSA to fuse EEG and fMRI, perceptual processing of complex visual scenes was found to start in early visual cortex ~100 ms post image onset, followed by ventral visual cortex at ~160 ms, in general agreement with prior reports.

### Formation of affect-specific neural representations

The question of how long it takes for affect-specific neural representations to form has been considered in the past. An intracranial electroencephalography study reported enhancement of gamma oscillations for emotional pictures compared to neutral pictures in occipital-temporal lobe in the time period of 200 ms −1000 ms (Boucher et al., 2015). In our data, the ~200 ms onset of above-chance decoding and ~500 ms occurrence of peak decoding accuracy, with the main contribution to decoding performance coming from occipital and temporal electrodes, are consistent with the previous report. Compared to nonaffective images such as faces, houses and scenes, where decodable differences in neural representations in visual cortex started to emerge ~100 ms post stimulus onset with peak decoding accuracy occurring at ~150 ms (Cichy et al., 2016; Cauchoix et al., 2014), these affect-specific times appear to be quite late. From a theoretical point of view, this delay may be explained by the reentry hypothesis which holds that anterior emotion regions such as the amygdala and the prefrontal cortex, upon receiving sensory input, send feedback signals to visual cortex to enhance sensory processing and facilitate motivated attention (Lang and Bradley, 2010). In a recent fMRI study (Bo et al., 2021), we found that affect-specific scenes can be decoded from multivoxel patterns in the retinotopic visual cortex and the decoding accuracy is correlated with the effective connectivity from anterior regions to visual cortex, in agreement with the hypothesis. What has not been established is how long it takes for the reentry signals to reach visual cortex. Providing a reference time for comparison by fusing EEG and fMRI data via RSA, we established that sensory processing of complex visual scenes such as those contained IAPS pictures began ~100 ms post picture onset, and thereby estimated that the reentry time is on the order of ~100 ms or shorter.

The negativity bias idea, which holds that negative events elicit more rapid and stronger responses compared to pleasant events (Rozin and Royzman 2001; Vaish et al., 2008), would seem to predict that neural representations of unpleasant images would form sooner than pleasant ones. While the idea has received support in behavioral data, e.g., subjects tend to locate unpleasant faces among pleasant distractors in shorter time than the reverse (Öhman et al., 2001), the neurophysiological support is mixed. Some studies using affective picture viewing paradigms reported shorter ERP latency and larger ERP amplitude for unpleasant pictures compared to pleasant ones in central P2 and late positive potential (LPP) (Carretié et al., 2001; Huang and Luo, 2006), but other ERP studies found that positive scene processing can be as strong and as fast as negative scene processing when examining early posterior negativity (EPN) in occipital channels (Schupp et al., 2006; Franken et al., 2008; Weinberg and Hajcak 2010). One possible explanation for the discrepancy is the choice of stimuli. The inclusion of exciting and sports images, which have high valence but low arousal, as stimuli in the pleasant category weakens the pleasant ERP effects when compared against threatening scenes included in the unpleasant category which have both low valence and high arousal (Weinberg and Hajcak 2010). Another possible contributing factor is the univariate nature of ERP analysis, which makes it mainly sensitive to effects in the localized regions but not those stemming from multiple concurrent neural processes taking place in distributed brain networks and their interactions. In the present work, by including images such as erotica and affiliative happy scenes in the pleasant category, which have comparable arousal ratings as images included in the unpleasant category, we were able to mitigate the possible issues associated with stimulus selection. In addition, by applying multivariate pattern analysis to EEG data from all channels, we were able to assess network contributions to the processing of affective scenes. Using above-chance level decoding as the indicator of the formation of emotion-specific representation, we presented evidence that is not consistent with the negative bias idea. In particular, when decoding between images from broad semantic categories (pleasant, unpleasant, and neutral), our results actually uncovered a positivity bias, where the neural representations of pleasant images formed earlier than that of unpleasant images. Subdividing the images into 6 subcategories: erotic couples, happy people, mutilation body/disgust, attack scene, neutral scene, and neutral people, and decoding the emotion subcategories against the neutral subcategories, we found the following temporal sequence of formation of neural representations: erotic couple (pleasant) → attack (unpleasant)→mutilation body/disgust (unpleasant), with happy people failing to be decoded from neutral images. This finding can be seen as providing neural support to previous electrodermal findings showing that erotic scenes evoked largest responses within IAPS pictures, which was followed by mutilation and threat scenes (Sarlo et al., 2005), suggesting the temporal dynamic of emotion processing depends on specific scene content.

### Temporal evolution of neural representations of affective scenes

Once the emotion-specific neural representations form, how do these representations evolve over time? If emotion processing is sequential, namely, if it progresses from one brain region to the next as time passes, we would expect a dynamically evolving neural pattern. On the other hand, if the emotional state is stable over time via recurrent processing in distributed brain networks, we would expect a sustained neural pattern. A technique for testing these possibilities is the temporal generalization method (King and Dehaene, 2014). In this method, a classifier trained on data at one time is applied to decode data from all other times, resulting in a 2D plot of decoding accuracy called the temporal generalization matrix. Past studies decoding between non-emotional images such as faces vs objects have found a transient temporal generalization pattern (Carlson et al., 2013; Cichy et al., 2014; Kaiser et al., 2016), supporting a sequential processing model for object recognition (Carlson et al., 2013). The temporal generalization from our data revealed that the neural representations of affective scenes are stable over a wide time window (~ 200 ms to 2000 ms), suggesting that the emotional state is maintained by sustained motivational attention, triggered by affective content (Schupp et al.,2004; Hajcak et al., 2009), which is in turn supported by recurrent interactions between sensory cortex and anterior emotion structures (Keil et al., 2009; Sabatinelli et al., 2009; Lang and Bradley 2010). The time window is broadly consistent with previous ERP studies where elevated LPP lasted multiple seconds and even extending beyond the offset of the stimuli (Foti and Hajcak, 2008; Hajcak et al., 2009).

### Role of visual cortex in sustained neural representations of affective scenes

The visual cortex, in addition to its role in processing perceptual information, is also expected to play an active role in sustaining affective representations, because the purpose of sustained motivational attention is to enhance vigilance towards threats or opportunities in the visual environment (Lang and Bradley, 2010). Sensory cortex’s role in sustained neural computations has been shown in other cognitive paradigms, including decision-making (Mostert et al., 2015), where stable neural representations are supported by the reciprocal interactions between prefrontal decision structures and sensory cortex. In face perception and imagery, neural representations are also found to be stable and sustained by communications between high and low order visual cortices (Dijkstra et al., 2018). In our data, two lines of evidence support a sustained role of visual cortex in emotion representation. First, over an extended time period, the weight maps obtained from EEG classifiers were comprised of channels located mainly in occipital-temporal areas. Second, if the emotion-specific neural representations in the visual cortex stem from the recurrent processing within distributed networks, then the stronger and longer these interactions, the stronger and more distinct the affective representations in visual cortex. This is supported by the finding that the strength of temporal generalization is correlated with the fMRI decoding accuracy in visual cortex.

### Temporal dynamic of sensory processing in visual pathway

The temporal dynamics of sensory processing of complex visual scenes can be revealed by fusing EEG-fMRI using RSA. The results showed that visual processing of IAPS images started ~100 ms post picture onset in early visual cortex and proceeded to ventral visual cortex at ~160 ms. This timing information is consistent with a previous ERP study where it is found that during the recognition of natural scenes, the low-level features are best explained by the ERP component occurring ~90 ms post picture onset while high-level features are best represented by the ERP component occurring ~170 ms after picture onset (Greene and Hansen, 2021). Compared with this time, the ~200 ms onset of affect-specific neural representations likely includes the time it took for the reentry signals to travel from emotion processing structures such as the amygdala or the prefrontal cortex to the visual cortex, which then give rise to the affect-specific representations seen on the scalp. The dorsal visual cortex, a brain region important for action and movement preparation (Wandell et al., 2011), is activated at ~360 ms, which is relatively late and may reflect the processing of action predispositions resulting from affective perceptions. This sequence of temporal activity is consistent with that established previously using the fast-fMRI method where early visual cortex activation preceded ventral visual cortex activation which preceded dorsal visual cortex activation (Sabatinelli et al., 2014). Additionally, the RSA similarity time courses in all three visual ROIs stayed highly activated for a relatively long time period, which may be taken as further evidence, along with the temporal generalization analysis, to support sustained representations of affective scenes. From a methodological point of view, the RSA differs from the decoding analysis in that decoding analysis captures affect-specific distinction between neural representations, whereas the RSA fusing of EEG-fMRI is sensitive to evoked pattern similarity shared by EEG and fMRI imaging modalities, with early effects likely driven by sensory perceptual processing and late effects by both sensory and affective processing.

## Materials and Methods

### Participants

The experimental protocol was approved by the Institutional Review Board of the University of Florida. Healthy volunteers (n=26) with normal or corrected-to-normal vision signed informed consent and participated in the experiment. Two participants withdraw before recording. Four additional participants were excluded for excessive movements inside the scanner. Data from the remaining 20 subjects were analyzed and reported here (10 women; mean age: 20.4±3.1). These data had been published before (Bo et al., 2021) but a different set of questions were addressed here.

### Procedure

#### The stimuli

The stimuli included 20 pleasant, 20 neutral and 20 unpleasant pictures from the International Affective Picture System (IAPS; Lang et al., 1997): Pleasant: 4311, 4599, 4610, 4624, 4626, 4641, 4658, 4680, 4694, 4695, 2057, 2332, 2345, 8186, 8250, 2655, 4597, 4668, 4693, 8030; Neutral: 2398, 2032, 2036, 2037, 2102, 2191, 2305, 2374, 2377, 2411, 2499, 2635, 2347, 5600, 5700, 5781, 5814, 5900, 8034, 2387; Unpleasant: 1114, 1120, 1205, 1220, 1271, 1300, 1302, 1931, 3030, 3051, 3150, 6230, 6550, 9008, 9181, 9253, 9420, 9571, 3000, 3069. The pleasant pictures included sports scenes, romance, and erotic couples and had average arousal and valence ratings of 5.8±0.9 and 7.0±0.5 respectively. The unpleasant pictures included threat/attack scenes and bodily mutilations and had average arousal and valence ratings of 6.2±0.8 and 2.8±0.8 respectively. The neutral pictures were images containing landscapes and neutral humans and had average arousal and valence ratings of 4.2±1.0 and 6.3±1.0 respectively. The arousal ratings for pleasant and unpleasant pictures are not significantly different (p=0.2); both are significantly higher than that of the neutral pictures (p<0.001). Based on specific content, the 60 pictures can be further divided into 6 subcategories: disgust/mutilation body, attack/threat scene, erotic couple, happy people, neutral people, and adventure/nature scene. These subcategories provided an opportunity to examine the content-specificity of temporal processing of affective images.

#### The paradigm

The experimental paradigm was illustrated in Figure 1A. There were five sessions. Each session contains 60 trials corresponding to the presentation of 60 different pictures. The order of picture presentation was randomized across sessions. Each IAPS picture was presented on a MR-compatible monitor for 3 seconds, followed by a variable (2800 ms or 4300 ms) interstimulus interval. The subjects viewed the pictures via a reflective mirror placed inside the scanner. They were instructed to maintain fixation on the center of the screen. After the experiment, participants rated the hedonic valence and emotional arousal level of 12 representative pictures (4 pictures for each broad category), which are not part of the 60-picture set, based on the paper and pencil version of the self-assessment manikin (Bradley and Lang, 1994).

### Data acquisition

#### EEG data acquisition

EEG data were recorded simultaneously with fMRI using a 32 channel MR-compatible EEG system (Brain Products GmbH). Thirty-one sintered Ag/AgCl electrodes were placed on the scalp according to the 10-20 system with the FCz electrode serving as the reference. An additional electrode was placed on subject’s upper back to monitor electrocardiogram (ECG); the ECG data was used during data preprocessing to assist in the removal of the cardioballistic artifacts. EEG signal was recorded with an online 0.1-250Hz band-pass filter and digitized to 16-bit at a sampling rate of 5 kHz. To ensure the successful removal of the gradient artifacts in subsequent analyses, the EEG recording system was synchronized with the scanner’s internal clock throughout recording.

#### fMRI data acquisition

Functional MRI data were collected on a 3T Philips Achieva scanner (Philips Medical Systems). The recording parameters are as follows: echo time (TE), 30 ms; repetition time (TR), 1.98 s; flip angle, 80°; slice number, 36; field of view, 224 mm; voxel size, 3.5*3.5*3.5 mm; matrix size, 64*64. Slices were acquired in ascending order and oriented parallel to the plane connecting the anterior and posterior commissure. T1-weighted high-resolution structural image was also obtained.

### Data preprocessing

#### EEG data preprocessing

The EEG data was first preprocessed using Brain Vision Analyzer 2.0 (Brain Products GmbH, Germany) to remove gradient and cardioballistic artifacts. To remove gradient artifacts, an artifact template was created by segmenting and averaging the data according to the onset of each volume and subtracted from the raw EEG data (Allen et al., 2000). To remove cardioballistic artifacts, ECG signal was low-pass-filtered, and the R peaks were detected as heart-beat events (Allen et al., 1998). A delayed average artifact template over 21 consecutive heart-beat events was constructed using a sliding-window approach and subtracted from the original signal. After gradient and cardioballistic artifacts were removed, the EEG data were lowpass filtered with the cutoff set at 50 Hz, downsampled to 250 Hz, re-referenced to the average reference, and exported to EEGLAB (Delorme and Makeig, 2004) for further analysis. The second-order blind identification (SOBI) procedure (Belouchrani et al., 1993) was performed to further correct for eye blinking, residual cardioballistic artifacts, and movement-related artifacts. The artifact-corrected data were then lowpass filtered at 30Hz and epoched from −300ms to 2000ms with 0ms denoting picture onset. The prestimulus baseline was defined to be −300ms to 0ms.

#### fMRI data preprocessing

The fMRI data were preprocessed using SPM (http://www.fil.ion.ucl.ac.uk/spm/). The first five volumes from each session were discarded to eliminate transient activity. Slice timing was corrected using interpolation to account for differences in slice acquisition time. The images were then corrected for head movements by spatially realigning them to the sixth image of each session, normalized and registered to the Montreal Neurological Institute (MNI) template, and resampled to a spatial resolution of 3mm by 3mm by 3mm. The transformed images were smoothed by a Gaussian filter with a full width at half maximum of 8 mm. The low frequency temporal drifts were removed from the functional images by applying a high-pass filter with a cutoff frequency of 1/128 Hz.

### MVPA analysis: EEG data

#### EEG decoding

To reduce noise and increase decoding robustness, 5 consecutive EEG data points (no overlap) were averaged, resulting in a smoothed EEG time series with a temporal resolution of 20 ms (50 Hz). Unpleasant vs neutral scenes and pleasant vs neutral scenes were decoded within each subject. The MVPA decoding was done at each time point to form a decoding accuracy time series. The 31 EEG channels provided 31 features for the SVM classifier. A ten-fold cross validation approach was applied. The weight vector from the classifier was transformed according to Haufe et al. (2014) and its absolute value is visualized as a topographical map to assess the discrepancy between affective and neutral state in each channel.

#### Temporal generalization

The stability of the neural representations evoked by affective scenes was tested using a generalization across time (GAT) method (King and Dehaene, 2014). In this method, the classifier was not only tested on the data from the same time point at which it was trained, it was also tested on data from all other sample points, yielding a two-dimensional temporal generalization matrix. The decoding accuracy at a point on this plane (*t_x_*, *t_y_*) reflects the decoding performance at time *t_x_* of the classifier trained at time *t_y_*.

#### Statistical significance testing of EEG decoding

Whether the decoding accuracy was above chance was evaluated by the Wilcoxon sign-rank test. Specifically, the decoding accuracy at each time point was tested against 50% (chance level). The resulting p value was corrected for multiple comparisons by controlling for the false discovery rate (FDR, p<0.05) across the time course. A further requirement to reduce possible false positives is that the significance cluster contains at least four consecutive such sample points.

The decoding accuracy was expected to be at chance level prior to and immediately after picture onset. The time at which decoding accuracy rose above chance level was taken to be the time when the neural representations of affective scenes formed. The statistical significance of the difference between the onset times of above-chance-decoding for different decoding accuracy time series was evaluated by a bootstrap resample procedure. Each resample consisted of randomly picking 20 sample decoding accuracy time series from 20 subjects with replacement and above-chance decoding onset was determined for this resample. The procedure was repeated 1000 times and the onset times from all the resamples formed a distribution. The significant difference between two such distributions was assessed by the two-sample Kolmogorov-Smirnov test.

### MVPA analysis: fMRI data

The picture-evoked BOLD activation was estimated on a trial-by-trial basis using the beta series method (Mumford et al., 2012). In this method, the trial of interest was represented by a regressor, and all the other trials were represented by another regressor. Six motion regressors were also included to account for any movement-related artifacts during scan. Repeating the process for all the trials we obtained the BOLD response to each picture presentation in all brain voxels. The single-trial voxel patterns evoked by pleasant, unpleasant, and neutral pictures were decoded between pleasant and neutral as well as between unpleasant and neutral using a ten-fold validation procedure within the retinotopic visual cortex defined according to a recently published probabilistic visual retinotopic atlas (Wang et al., 2014). Here the retinotopic visual cortex consisted of V1v, V1d, V2v, V2d, V3v, V3d, V3a, V3b, hV4, hMT, VO1, VO2, PHC1, PHC2, LO1, LO2, and IPS. For some analyses, the voxels in all these regions were combined to form a single ROI called visual cortex, whereas for other analyses, these regions were divided into early, ventral, and dorsal visual cortex (see below).

### Fusing EEG and fMRI data via RSA

Decoding between different emotion categories yields information on the formation and dynamics of affect-specific neural representations. For comparison, we also obtained the timing of perceptual or sensory processing of affective images in visual cortex by fusing EEG and fMRI data via representation similarity analysis (RSA) (Kriegeskorte et al., 2008). Trials from all emotion categories were combined so that the results would be about perceptual or sensory processing rather than affect-specific processing. For EEG data, 31 channels of EEG data at a given time point provided a 31-dimensional feature vector. The 300 × 300 representational dissimilarity matrix (RDM) was constructed by pairwise correlating the feature vectors from all 300 trials (60 trials per session x 5 sessions). For fMRI data, following previous work (Bo et al., 2021), we divided the visual cortex into three ROIs: early (V1v, V1d, V2v, V2d, V3v, V3d), ventral (VO1, VO2, PHC1, PHC2), and dorsal (IPS1-5) visual cortex. The fMRI feature vector was extracted from single-trial beta-series within each ROI. A 300 × 300 RDM was constructed for each ROI by pairwise correlating the 300 feature vectors from 300 trials. The RDM from EEG data at each time point and the RDM from fMRI data in each ROI were correlated, yielding a representational similarity time course for each ROI, with the similarity at a given time point reflecting the common variance captured by simultaneous EEG and fMRI for that ROI at that time point.

To assess the onset time of significant similarity between EEG RDM and fMRI RDM, we first computed the mean and standard deviation of the similarity measure during the baseline period (−300 ms to 0 ms). The mean and standard deviation is averaged across ROIs. Along the representational similarity time course, similarity measures that are five standard deviations above the baseline mean were considered statistically significant (p<0.003). To further control for multiple comparisons, clusters containing smaller than five consecutive such time points were discarded. For a given ROI, the first time point that meets the above significance criteria was considered the onset time for perceptual or sensory processing for that ROI. To compare the three onset times from the three ROIs, we conducted a bootstrap resample procedure. Each resample consisted of randomly picking 20 sample RDM similarity time series from the 20 subjects with replacement and the onset time was determined for the resample. The procedure was repeated 1000 times and the onset times from all the resamples formed a distribution. The significant difference between distributions was then assessed by the two-sample Kolmogorov-Smirnov test.

## Acknowledgements

This work was supported by NIH grants R01 MH112558 and R01 MH125615.

## Notes

### Competing Interest Statement

The authors have declared no competing interest.

